# CH-related Mutant ASXL1 Promotes Atherosclerosis in Mice via Dysregulated Innate Immunity

**DOI:** 10.1101/2024.08.18.608516

**Authors:** Naru Sato, Susumu Goyama, Yu-Hsuan Chang, Takeshi Fujino, Tamami Denda, Shuhei Asada, Shiori Shikata, Xiaoxiao Liu, Taishi Yonezawa, Reina Takeda, Keita Yamamoto, Yosuke Tanaka, Hiroaki Honda, Yasunori Ota, Takuma Shibata, Motohiro Sekiya, Shuhei Koide, Chrystelle Lamagna, Esteban Masuda, Atsushi Iwama, Hitoshi Shimano, Jun-ichiro Inoue, Kensuke Miyake, Toshio Kitamura

## Abstract

Certain somatic mutations confer a fitness advantage in hematopoietic stem and progenitor cells (HSPCs) over normal HSPCs, resulting in the clonal expansion of mutant blood cells^1^, otherwise known as clonal haematopoiesis (CH). CH is frequently observed among healthy elderly people and is closely associated with the risk of cardiovascular diseases (CVDs). The most frequently mutated genes of CH include *DNMT3A, TET2*, and *ASXL1*^2^. Among them, even though *ASXL1* mutations are clinically associated with the highest risk for developing CVDs, little is known whether and how the mutations contribute to CVDs. Here we show accelerated development of atherosclerosis and increased inflammatory monocytes in mice transplanted with the bone marrow cells (BMCs) from the mice expressing mutant ASXL1 (ASXL1-MT) selectively in hematopoietic cells. RNA sequencing analysis of the plaque-macrophages derived from BMCs expressing ASXL1-MT showed more inflammatory signatures than those from control BMCs. Mechanistically, wild-type ASXL1 inhibited innate immune signalling through direct interactions with IRAK1/TRAF6/TAK1 in the cytoplasm, while ASXL1-MT, which only interacted with TAK1, lost this regulatory function, leading to NF-κB activation. This mechanism is unique and distinct from those of CH with *Tet2* or *Dnmt3a* mutations, where overactivation of the IL-1β/NLRP3 inflammasome plays critical roles^3–5^. Intriguingly, IRAK1/4 inhibition decreased the number of inflammatory monocytes and attenuated the development of atherosclerosis driven by ASXL1-MT. The present work connects the mutations of an epigenetic factor, ASXL1, with inflammation and CVDs and gives an indication for the prevention of CVDs in CH.

---

Somatic mutations accumulate in hematopoietic stem and progenitor cells (HSPCs) as we age. Certain mutations confer a fitness advantage in these HSPCs over normal HSPCs without mutations, resulting in the clonal expansion of mutant blood cells ^1^. Clonal hematopoiesis (CH) is characterized by the clonal expansion of mutant blood cells in persons without other hematologic abnormalities including anemia. CH is common among the elderly and is associated with an increased risk of hematopoietic neoplasms and mortality ^6–8^.

Epidemiological studies have revealed a significant association between CH and cardiovascular diseases (CVDs)^9^, and several reports have suggested the causal effect of CH on CVDs using mouse models ^10^. Tet2 deficiency in bone marrow cells (BMCs) led to an increase of NLRP3 inflammasome-mediated IL-1β secretion in macrophages and a marked increase in atherosclerotic plaque size in mice ^3,4^. On the other hand, loss of Dnmt3a in BMCs worsened heart function through greater macrophage accumulation and increased expression of immune cell markers in myocardium^11^, suggesting that this condition impairs the resolution of inflammation overactivation of the IL-1β/NLRP3 inflammasome^5^.

Although clinical observations have indicated a close link between *ASXL1* CH and CVDs, the roles of *ASXL1*-mutated blood cells in the development of atherosclerosis and CVDs have not been experimentally investigated. ASXL1 plays key roles in epigenetic regulation in cooperation with multiple chromosome modifiers in the nucleus, and its mutations always predict a poor prognosis of haematological malignancies^12^. Most *ASXL1* mutations are nonsense or frameshift mutations of the last exon, resulting in the expression of C-terminally truncated mutant proteins^13^. Previously, we generated a conditional knockin (KI) mouse as a model for CH with the hematopoietic lineage-specific expression of mutant ASXL1 (ASXL1-MT)^14^. The present study provides a unique mechanism by which ASXL1-MT promotes atherosclerosis.

## Results

### High-fat diet and ASXL1-MT collaborate in inducing the expansion of inflammatory monocytes

Diabetes Mellitus (DM) is one of the conventional risk factors of CVDs^15^. Because the association of CH and DM has been reported^16^, we first assessed the effect of *ASXL1* mutation in haematopoietic cells on the glucose tolerance. ASXL1-MT KI mice (8-weeks old) presented no symptoms related to DM (Extended Data Fig. 1a). We then fed control and ASXL1-MT KI mice a high-fat diet (HFD) or normal chow for 10 weeks. HFD increased body weight and reduced the glucose tolerance in both control and ASXL1-MT KI mice equally (Extended Data Fig. 1a-e). As for blood parameters, the number of white blood cells and inflammatory monocytes increased in ASXL1-MT KI mice on HFD (Extended Data Fig.1f). The numbers of neutrophils and patrolling monocytes tended to increase in HFD-fed ASXL1-MT KI mice. These results indicate that HFD and ASXL1-MT synergistically induce the expansion of inflammatory monocytes, while haematopoiesis with ASXL1-MT doesn’t affect the glucose tolerance nor obesity.

We next assessed the influence of HFD on the in vivo growth of bone marrow (BM) cells expressing ASXL1-MT using the competitive repopulation assay (Fig. 1a). As reported previously^14,17^, ASXL1-MT KI cells showed an impaired repopulation (Extended Data Fig. 2a) but a quicker recovery four months after the transplantation when fed the HFD (Fig. 1b). Similarly, neutrophils and Ly6C^hi^ inflammatory monocytes were increased among ASXL1-MT KI cells in recipient mice on HFD (Fig. 1c, Extended Data Fig. 2b).

**Fig. 1.**
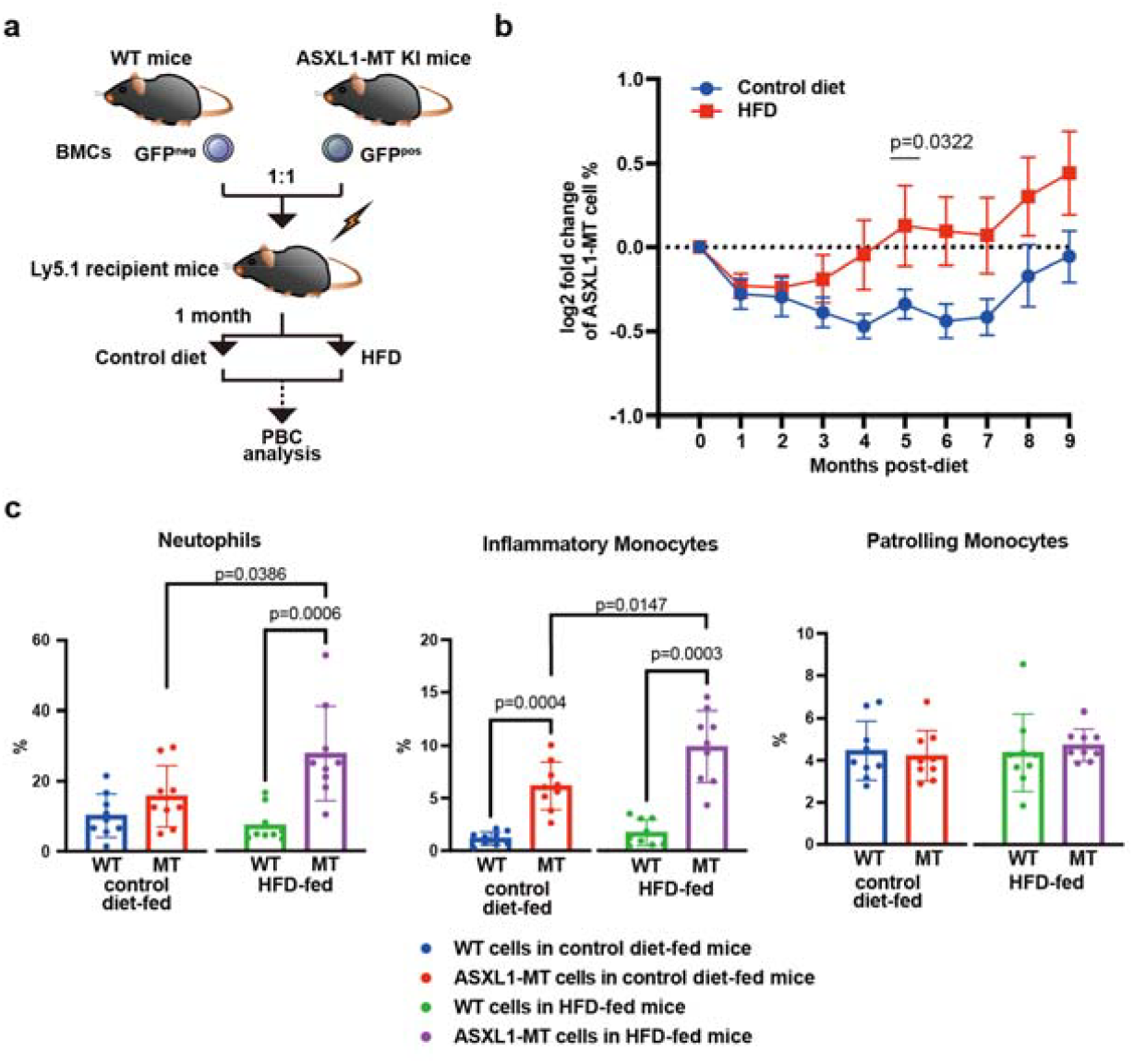
HFD confers a growth advantage on ASXL1-MT cells, particularly in the population of inflammatory monocytes. a: The experimental design for competitive transplantation assays with high-fat diet (HFD) feeding. Briefly, bone marrow cells (BMCs) from *Vav-cre* ASXL1-MT KI mice and littermate control mice were mixed at a 1:1 ratio, followed by transplantation into lethally irradiated Ly5.2 recipient mice. Four weeks later to allow for hematopoietic reconstitution, mice were separated into a HFD-fed group and control diet-fed group. b: Fold change of the chimerism of GFP+ ASXL1-MT cells in peripheral blood compared with the chimerism at the start of HFD or control diet feeding. (n = 9 (5 female and 4 male) per each group, respectively). Data are the mean ± s.e.m. Unpaired t-tests were used for the statistical analysis. c: The frequencies of neutrophils, inflammatory monocytes, and patrolling monocytes in peripheral blood cells of GFP-control or GFP+ ASXL1-MT cells in mice fed a control diet or HFD at 6 months after transplantation (n = 9 for each group). The percentage of neutrophils, inflammatory monocytes, and patrolling monocytes derived from ASXL1-MT KI bone marrow (MT) or littermate-control bone marrow (WT) in control diet-fed mice or HFD-fed mice in the competitive transplantation assay after 5-months of feeding. (n = 7 mice per group). See also Extended Data Fig. S1. Data are the mean ± s.d. Paired t-tests (c: comparison between MT and WT) and unpaired t-tests (b and c: comparison between HFD-fed mice and control diet-fed mice) were used for the statistical analysis.

### ASXL1-MT induces IL-6 production in macrophages and accelerates atherosclerosis in Ldlr knockout (*Ldlr*^-/-^) mice

To assess the role of *ASXL1* CH on atherosclerosis, we transplanted BM cells from control or ASXL1-MT KI mice (crossed with *Vav-cre* mice) into lethally irradiated atherosclerosis-prone low-density lipoprotein receptor knockout (*Ldlr*^-/-^) mice (Fig. 2a). These mice were fed a high fat/high cholesterol diet (HFHCD) for 12 weeks to induce CVD. Both female (Fig. 2b, c, Extended Data Fig. 3a, b) and male (Extended Data Fig. 3c, d) mice reconstituted with ASXL1-MT KI cells showed larger atherosclerotic lesions, but no differences in body weight or sizes of the liver, spleen, or kidney (Extended Data Fig. 3e, f). Masson trichrome staining showed thin fibrous caps of the atherosclerotic lesions (Extended Data Fig. 3g, h), which are signs for increased myocardial infarction risk ^18^. The frequencies of neutrophils and monocytes increased in the peripheral blood of *Ldlr*^*-*/-^ mice that received ASXL1-MT KI BM cells (Extended Data Fig. 3i). Next, we crossed ASXL1-MT KI mice with *LysM-cre* mice to limit ASXL1-MT expression in myeloid cells (Extended Data Fig. 4a, b). BM cells derived from control or *LysM-cre* ASXL1-MT KI mice were then transplanted into lethally irradiated *Ldlr*^-/-^ mice, which were fed the HFHCD for 14 weeks after the hematopoietic reconstitution. As with *Vav-cre*/ASXL1-MT KI mice, we observed an increased plaque size in the aortic root of transplanted *Ldlr*^-/-^ mice (Extended Data Fig. 4c), indicating that myeloid cells play major roles in atherosclerosis.

**Fig. 2.**
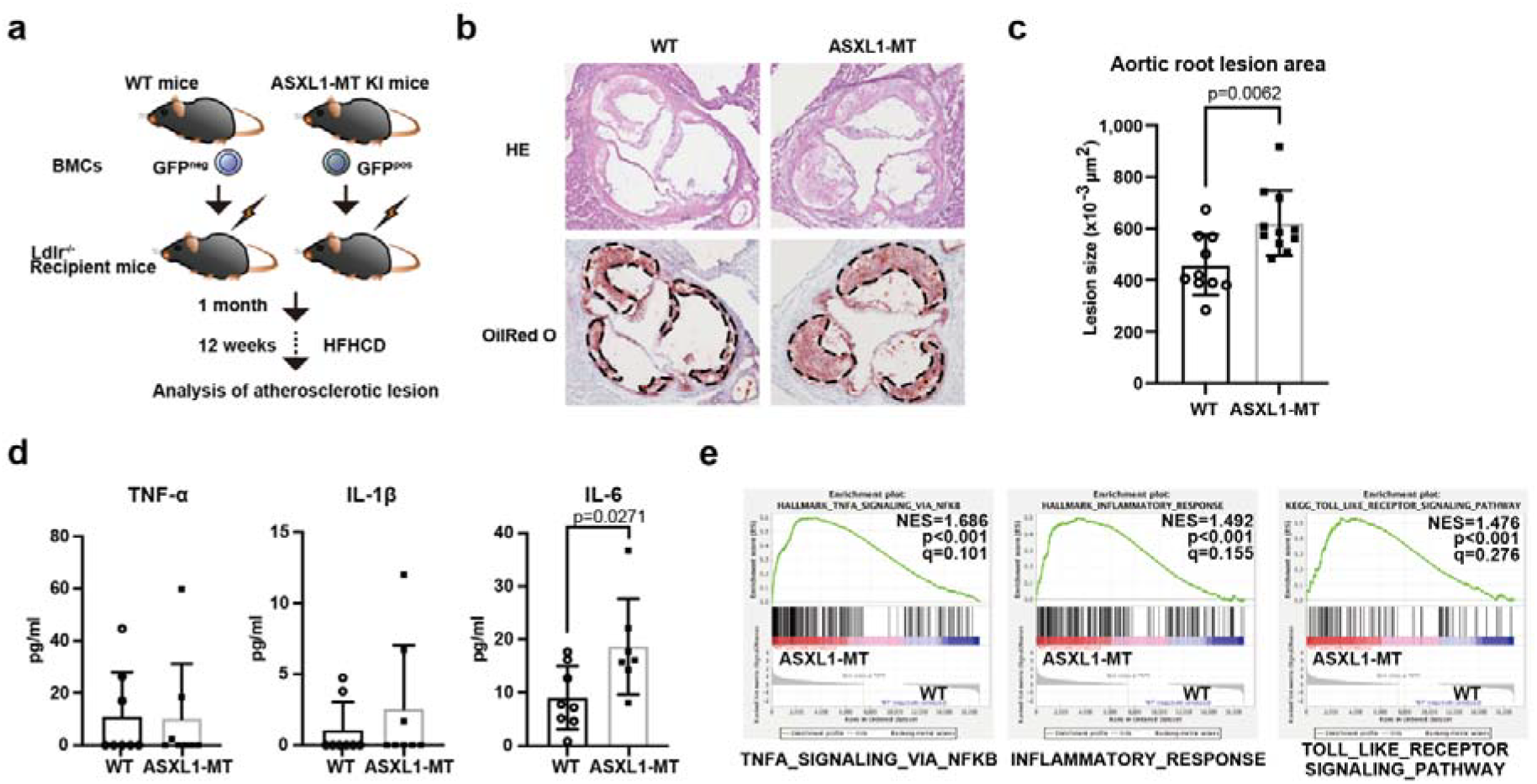
Mutant ASXL1-derived hematopoiesis promotes atherosclerosis in *Ldlr*^-/-^ mice. a: Scheme of the experimental procedures for atherosclerosis studies in mice with hematopoietic-specific expression of ASXL1-MT. Briefly, bone marrow cells (BMCs) from *Vav-cre* ASXL1-MT KI mice or littermate control mice were transplanted into lethally irradiated *Ldlr*^-/-^ recipient mice. Four weeks after the transplantation, mice were fed a high-fat/high-cholesterol diet (HFHCD) for 12 weeks (female) or 14 weeks (male). b: Representative images of aortic root lesions that were stained with H&E (upper) or Oil Red-O (lower) from female *Ldlr*^-/-^ mice with WT or *Vav-cre* ASXL1-MT KI bone marrow after 12-weeks of HFHCD feeding. The dashed lines indicate the atherosclerotic lesion areas. c Quantification of the lesion area in the aortic root (n= 10 (mice transplanted with control BMCs) and 11 (mice transplanted with ASXL1-MT KI BMCs)). d: Serum levels of Tnf-α, Il-6, and Il-1β in *Ldlr*^-/-^ mice (n = 8 control, n = 7 ASXL1-MT, respectively). e: Gene set enrichment analysis (GSEA) of aortic macrophages showing gene sets positively or negatively associated with mice transplanted with ASXL1-MT BMCs or mice transplanted with control BMCs (n=3 per group).

To investigate the differentiation of HSPCs harbouring ASXL1-MT compared to normal HSPCs under HFHCD conditions, competitive transplantation was performed (Extended Data Fig. 4d). The frequencies of neutrophils and inflammatory monocytes but not patrolling monocytes derived from ASXL1-MT KI BM increased 20 weeks after the transplantation in HFHCD-fed *Ldlr*^-/-^ mice (Extended Data Fig. 4e, f), indicating that ASXL1-MT skewed the fate of HSCs to myeloid, particularly inflammatory monocytes and neutrophils, when fed on HFHCD.

Because atherosclerosis is caused by chronic inflammation, we examined the levels of major pro-inflammatory cytokines in the serum of HFHCD-fed *Ldlr*^-/-^ mice. Mice transplanted with ASXL1-MT KI cells showed significantly higher serum levels of IL-6 but not TNF-α and IL-1β than control mice (Fig. 2d). We then generated BM-derived macrophages (BMDMs) from ASXL1-MT KI and control mice and treated the cells with a Toll-like receptor 4 (TLR4) agonist, lipopolysaccharide (LPS). We observed a significant upregulation of IL-6, but not of TNF-α in LPS-treated ASXL1-MT KI BMDMs (Extended Data Fig. 5a). RNA-seq analysis and gene set enrichment analysis (GSEA) of the CD11b^+^ F4/80^+^ aortic macrophages derived from ASXL1-MT KI BM revealed the activation of inflammatory responses, TLR signalling and NF-κB (Fig. 2e). The up-regulation of NF-κB canonical downstream genes (Nlrp3, Tnfaip3, and Irak3) was observed in aortic macrophages derived from ASXL1-MT KI BM, but NIK (Map3k14), a hallmark of the noncanonical pathway^19^, was not upregulated (Extended Data Fig. 5b). Taken together, these data indicate that ASXL1-MT activates TLR signalling, leading to activation of the canonical NF-κB pathway and an increase of IL-6 production in myeloid cells, thereby accelerating the development of atherosclerosis in *Ldlr*^-/-^ mice.

### Cytoplasmic ASXL1 inhibits the innate immune pathway

We next used a reporter assay system for TLRs to clarify the potential roles of ASXL1 in the regulation of innate immune pathways. We transduced a plasmid harbouring wild-type ASXL1 (ASXL1-WT) or ASXL1-MT into Ba/F3 cells expressing an NF-κB luciferase reporter^20^ and the indicated TLRs (BaκB cells) and stimulated these reporter cells by the corresponding ligands (Fig. 3a). Interestingly, ASXL1-WT attenuated TLR5- and TLR7-induced NF-κB activation, while ASXL1-MT caused slightly enhanced them (Fig. 3b).

**Fig. 3.**
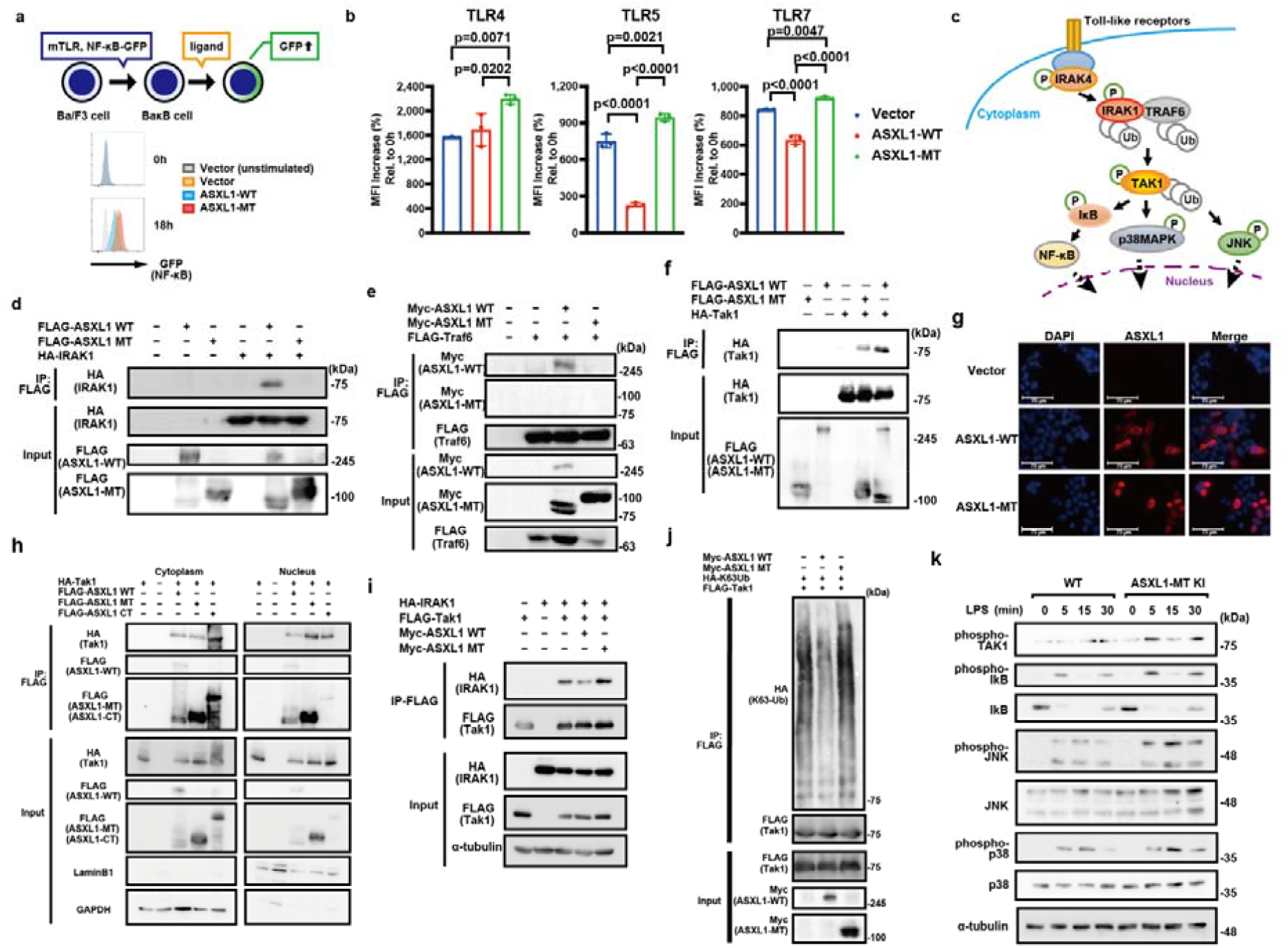
ASXL1-WT, but not ASXL1-MT, inhibits activation of TAK1 via direct interaction with the IRAK1-TRAF6 complex in the cytoplasm. a: The scheme and a representative histogram of Ba/F3 cells transduced with NF-κB GFP reporter gene (BaκB cells) expressing Toll-like receptor (TLR). BaκB cells were transduced with vector (mock) or ASXL1-WT or ASXL1-MT and sorted by flow cytometry. Colored histograms show the upregulated GFP expression in BaκB cells expressing TLR5 18 h after stimulation. The shaded histogram shows the background without stimulation. b: BaκB cells expressing TLR4 or TLR5 or TLR7 were stimulated by Lipid A (1μg/ml), flagellin (5 ng/ml) or R848 (100 ng/ml). Upregulated GFP expression by each stimulation was analyzed by flow cytometry. At 2 weeks and 4 weeks after sorting, cells were stimulated with R848 (100 ng/ml) for 18 h and GFP induction was assessed by flow cytometry. The results are represented by the % MFI increase normalized by that from the cells transduced with an empty vector (pMYs-IRES-hNGFR). Data are the mean ± s.d. and were assessed by one-way ANOVA with Tukey–Kramer’s post-hoc test. c: Overview of the TLR-NF-κB signaling pathway mediated by the activation of IRAK1, TRAF6, and TAK1. The activation of the signaling pathway is controlled by post-translational modifications of the signaling components, including phosphorylation and ubiquitination. A protein complex composed of IRAK1 and TRAF6 (E3) induces K63 polyubiquitination of TAK1. The K63-linked polyubiquitin chains do not induce proteasomal degradation but act as platforms for the activation of downstream molecules such as IκB, p38 MAPK, and JNK. d: 293T cells were transfected with 3xFLAG-ASXL1-WT or 3xFLAG-ASXL1-MT expression plasmid together with a plasmid encoding HA-IRAK1. Cell lysates were subjected to immunoprecipitation with an anti-FLAG antibody followed by immunoblotting with an anti-HA antibody. Whole-cell lysate (WCL) was also prepared and analyzed by immunoblotting with an anti-HA antibody and an anti-FLAG antibody. e: 293T cells were transfected with a Myc-ASXL1-WT or a Myc-ASXL1-MT expression plasmid together with a plasmid encoding FLAG-Traf6. WCL was immunoprecipitated with an anti-FLAG antibody, and Traf6-bound ASXL1-WT was detected with an anti-Myc antibody. WCL was also prepared and analyzed by immunoblotting with an anti-Myc antibody and an anti-FLAG antibody. f: 293T cells were transfected with 3xFLAG-ASXL1-WT or 3xFLAG-ASXL1-MT expression plasmid together with a plasmid encoding HA-Tak1. WCL was subjected to immunoprecipitation with an anti-FLAG antibody followed by immunoblotting with an anti-HA antibody. g: 293T cells were transfected with 3xFLAG-ASXL1. The cells were fixed and subjected to immunofluorescence staining. ASXL1 (red) is superimposed over nuclei stained with DAPI (blue). Representative images for Vector control (top), ASXL1-WT (middle), and ASXL1-MT (bottom) are shown. Scale bars: 75 μm. Images were captured by EVOS. See also Fig.3F. h: 293T cells were transfected with 3xFLAG-ASXL1-WT, 3xFLAG-ASXL1-MT or 3xFLAG-ASXL1-CT expression plasmid together with a plasmid encoding HA-Tak1. Nuclear (right) and cytoplasmic (left) protein fractions were separated using the hypotonic buffer and were subjected to immunoprecipitation with an anti-FLAG antibody followed by immunoblotting with an anti-HA antibody. See also Fig.3E. i: 293T cells transfected with Myc-ASXL1-WT, Myc-ASXL1-MT, Vector, FLAG-Tak1, or HA-IRAK1 and were immunoblotted for HA (IRAK1) on immunoprecipitated FLAG-Tak1. j: 293T cells transfected with Myc-ASXL1-WT, Myc-ASXL1-MT, FLAG-Tak1, or mutant ubiquitin (Ub) that forms only K63 linkages were immunoblotted for HA (K63Ub) on immunoprecipitated FLAG-Tak1. k: Immunoblot analysis of bone marrow-derived macrophages (BMDMs) from WT and *Vav-cre* ASXL1-MT KI bone marrow cells. BMDMs were stimulated with 1μg/ml LPS for the indicated times (in minutes) to evaluate the phosphorylation of Tak1, IκB, JNK, and p38 MAPK.

To investigate the underlying molecular mechanisms of the ASXL1-WT effects, we examined the physical interaction between ASXL1 and major components of the innate immune pathway in 293T cells (Fig. 3c, Extended Data Fig. 6a). ASXL1-WT bound to IRAK1, TRAF6, TAK1, TAB2, and TAK1-TAB3, while ASXL1-MT interacted only with TAK1 (Fig. 3d-f, Data Fig. 6b, c). Interestingly, we noticed that the co-expression of IRAK1, TRAF6, TAK1, TAB2, or TAB3 and ASXL1-WT increased the expression of a short form of ASXL1, which had a similar size as ASXL1-MT and displayed 100 kDa bands (Fig. 3d-f, Extended Data Fig. 6b-e). However, this short form was not detected when the catalytically inactive TAK1 (TAK1-K63W) was co-expressed with ASXL1-WT (Extended Data Fig. 6d). These results demonstrate that ASXL1-WT physically interacts with multiple components involved in TLR signalling and suggest that inflammatory signals could increase the amount of the short form of ASXL1, for which TAK1 activity is required.

We next examined the subcellular localization of ASXL1-WT and ASXL1-MT in 293T cells and found that ASXL1-WT existed and interacted with TAK1 and IRAK1 both in the nucleus and cytoplasm. In contrast, ASXL1-MT preferentially localized in the nucleus but interacted with TAK1 both in the nucleus and cytoplasm (Fig. 3g, h, Extended Data Fig. 6e). A C-terminal lesion of ASXL1 (ASXL1-CT), which ASXL1-MT lacks, also bound to TAK1 (Fig. 3h, Extended Data Fig. 6b), indicating that ASXL1 binds TAK1 via both its N-terminal and C-terminal regions. To avoid any artificial effects by the over-expressed proteins, we also examined the subcellular localization of endogenous ASXL1 in BMDMs and confirmed that ASXL1 mainly localizes in the cytoplasm but with a fraction expressed in the nucleus as well (Extended Data Fig. 6f). These data corroborate a previously unrecognized role of ASXL1-WT to suppress TAK1 activation.

TAK1 is a kinase that plays critical roles in the activation of the TLRs-NF-κB pathway. TAK1 is activated by IRAK-mediated phosphorylation and TRAF6-mediated lysine 63 (K63)-linked polyubiquitination resulting in recruiting many signalling molecules^21^. To further investigate how ASXL1-WT inhibits TAK1 activation, we examined TAK1 interaction with IRAK1. We identified that ASXL1-WT, but not ASXL1-MT, inhibited the interaction between IRAK1 and TAK1 and profoundly suppressed the K63-linked polyubiquitination of TAK1 in 293T cells (Fig. 3i-k). Both ASXL1-WT and ASXL1-MT bound with TAK1-K63W, a TAK1 mutant that does not incur K63-linked polyubiquitination^22^, suggesting that K63-linked polyubiquitination is not necessary for the binding between ASXL1 and TAK1 (Extended Data Fig. 6d). Moreover, we observed an increased phosphorylation of TAK1 as well as several downstream molecules, including p38MAPK, IκB, and JNK, upon LPS stimulation in BMDMs derived from ASXL1-MT KI mice when compared to those derived from control mice (Fig. 3k). These results indicate that cytoplasmic ASXL1 prevents IRAK1-TAK1 interaction and suppresses IRAK1/TRAF6-mediated TAK1 activation. Importantly, ASXL1-MT lacks this inhibitory effect.

### IRAK1/4 inhibitor suppresses atherosclerosis induced by ASXL1-MT KI blood cells

Finally, we examined whether inhibition of the innate immune pathway by an IRAK1/4 inhibitor suppresses the development of atherosclerosis (Fig. 4a). The IRAK1/4 inhibitor R191^23^ inhibited the ubiquitination of TAK1 as well as the phosphorylation of several downstream molecules, including IκB, JNK and p38MAPK (Extended Data Fig. 7a-c). We transplanted ASXL1-WT or ASXL1-MT KI BM cells into *Ldlr*^-/-^ mice, which were fed a control diet, HFHCD or HFHCD containing another IRAK1/4 inhibitor, R221, a prodrug of R191 suitable for *in vivo* administration (Extended Data Fig. 7d). A 6-week treatment with R221 reverted the ASXL1-MT-induced increase of neutrophils and inflammatory monocytes but had little effect on the number of myeloid cells derived from control BM cells (Fig. 4b). Moreover, a 12-week treatment with R221 significantly reduced the size of the aortic plaques in mice transplanted with ASXL1-MT KI BM cells (Fig. 4c, d). Intriguingly, this inhibitory effect of R221 on atherosclerosis appears to be specific to the ASXL1-MT KI model; the same treatment did not reduce the size of the aortic plaques in mice that received wild-type BM cells (Extended Data Fig. 7e, Extended Data Table 1). These results demonstrate that the aberrant myelopoiesis and exacerbated atherosclerosis induced by ASXL1-MT could be prevented by IRAK1/4 inhibitors and suggest that the mechanisms by which ASXL1-MT provides an atherosclerosis-prone state are different from that for conventional risks.

**Fig. 4.**
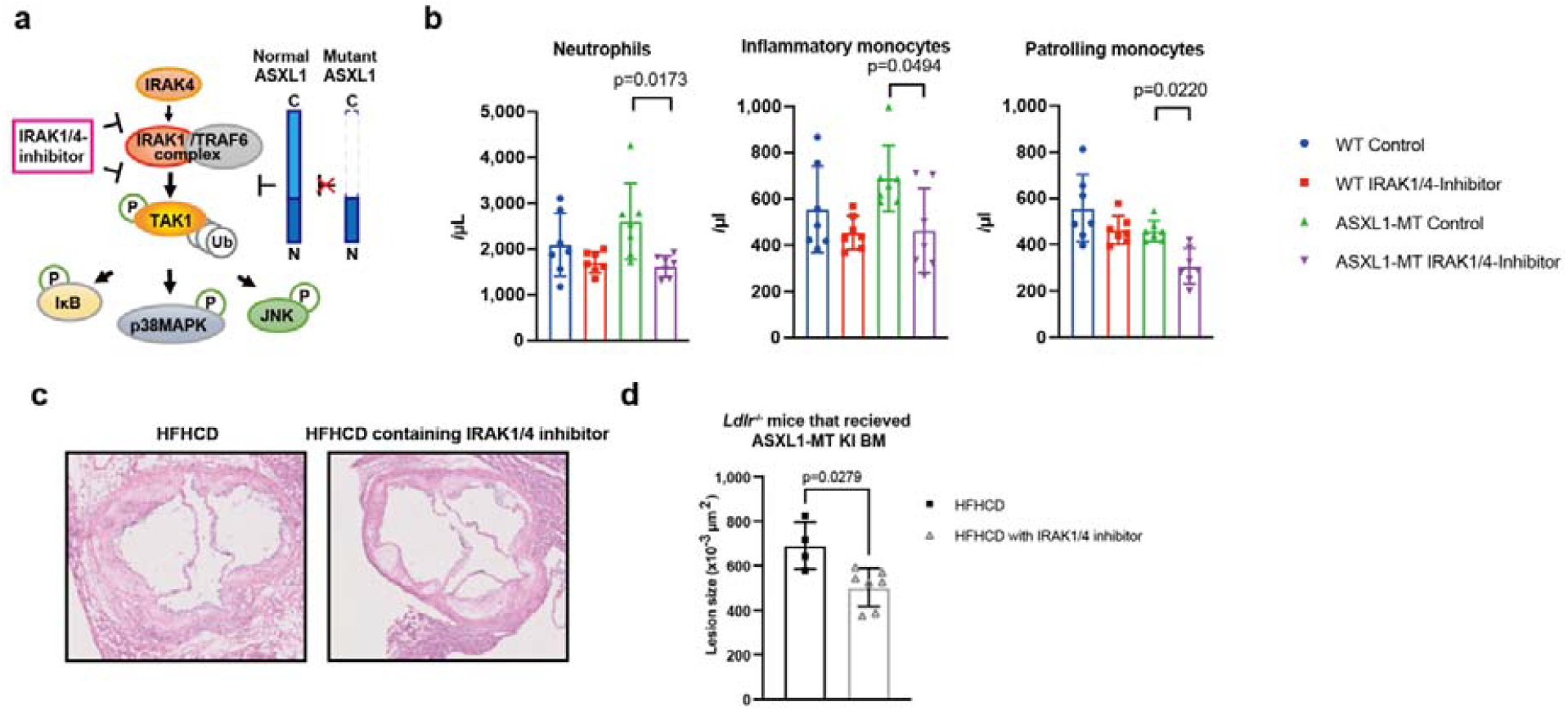
IRAK1/4 inhibitor improves mutant ASXL1-induced aberrant myelopoiesis and atherosclerosis. a: Overview of the TLR-mediated activation of NF-κB, and the proposed regulation by ASXL1-WT. ASXL1-WT binds to the IRAK1-TRAF6 complex to inhibit the activation of TAK1, while ASXL1-MT loses this regulatory function due to the inability to bind to the IRAK1-TRAF6 complex. IRAK1/4 inhibitor R191 inhibits the activation of TAK1. b: Summary of donor-derived PBC populations at 6 weeks after feeding with each diet. (n = 7 per group.) Colored horizontal lines represent median values. Data were assessed by one-way ANOVA with Tukey–Kramer’s post-hoc test. c, d: Representative images of the aortic root (c) and quantification of the lesion areas (d) that were stained with H&E from female *Ldlr*^-/-^ mice with *Vav-cre* ASXL1-MT KI BM after feeding. (n= 13 mice fed HFHCD, 7 mice fed R221-containing HFHCD). Data are presented as the mean ± s.d. Unpaired t-tests with Welch’s correction were used for the statistical analysis.

## Discussion

We here show the atheropromotive effect of ASXL1-MT. Based on epidemical information^24^ and our experimental results, the inflammatory environment associated with overeating or smoking may be closely related to the expansion of *ASXL1* mutated clones. These findings suggest the existence of a positive feedback loop between *ASXL1* CH and the inflammatory environment: ASXL1-MT induces aberrant myelopoiesis, which produces a pro-inflammatory environment and the inflammatory environment in turn promotes the expansion of *ASXL1* mutated clones (Extended Data Fig. 8). This vicious cycle is likely to underlie the development of the atherosclerosis associated with *ASXL1* CH. In fact, an increase of monocytes and serum C-reactive protein (CRP) levels were reported in CH carriers with *ASXL1* mutations^25^.

Mechanistically, our study revealed the unexpected cytoplasmic functions of ASXL1 as a modulator of the innate immune pathway. ASXL1 directly inhibits the IRAK1/TRAF6-induced activation of TAK1 and downstream signalling in association with the decreased K63-ubiquitination of TAK1, whereas ASXL1-MT lost this activity. Consistent with the present results, Muto et al. reported that the overexpression of TRAF6 observed in patients with myelodysplastic syndrome (MDS) contributes to the expansion of the cell clone via activation of the NF-kB pathway^26^. While they found that TRAF6 overexpression induced inflammation via the non-canonical NF-κB pathway, we show that ASXL1-MT induces inflammation mainly through the canonical NF-κB, implicating distinct signal transduction between the two conditions. While both TRAF6 overexpression and the expression of ASXL1-MT are frequently observed in patients with MDS, ASXL1 binds to downstream components of TLR and IL-1R, consistent with the fact that ASXL1-MT mainly activates the canonical NF-κB pathway.

Clinically, high concentrations of serum IL-6 are observed in human CH carriers including *ASXL1*-CH. In addition to IL-6, CH carriers with *TET2* mutations showed high levels of IL-1β, while *JAK2* and *SF3B1* mutations in blood cells were associated with increased circulating IL-18. We here demonstrate that ASXL1-MT activates or fails to inhibit the TLR signalling pathway, resulting in the activation of NF-κB, p38 MAPK, and JNK, leading to increased IL-6 production in macrophages and enhanced inflammation. In fact, high levels of CRP are detected only in CH harbouring *ASXL1* mutations, indicating the close association of inflammation and *ASXL1* mutations^25^.Thus, each CH-associated mutation appears to have distinct functions in the regulation of inflammatory responses.

Considering that the overexpression of IRAK1, TRAF6, or TAK1 increased the short forms of ASXL1 migrating near ASXL1-MT, it is tempting to speculate that inflammation somehow induces proteolytic products of ASXL1 that have a similar C-terminal truncation as ASXL1-MT and that inflammation becomes prolonged with the continuous expression of ASXL1-MT. To prove this hypothesis, further studies are required to identify which signal induces the proteolysis of ASXL1-WT and by which protease.

CH is an important issue in ageing societies. The Canakinumab Anti-inflammatory Thrombosis Outcomes Study (CANTOS) showed a reduction of cardiovascular events with an anti-IL-1β antibody, canakinumab^27^. Furthermore, previous studies have shown the atheroprotective activity of IL-1β blockade in mouse CH models for TET2 or JAK2 mutations harbouring dysregulated inflammasome-activation^4,28^. However, IL-1β-targeting therapy had no significant benefit for patients with IL-6 levels equal to or above 1.65 ng/l after taking the first dose of canakinumab^29^. This therapy is probably less effective at preventing atherosclerosis associated with *ASXL1* mutations, because the serum IL-6 level, but not the IL-1β level, is increased in human *ASXL1* CH carriers like in ASXL1-MT KI mice. Instead, we found that an IRAK1/4 inhibitor suppressed the aberrant myelopoiesis and reverted the exacerbated atherosclerosis in HFHCD-fed *Ldlr*^-/-^ mice transplanted with ASXL1-MT KI BM cells but not with normal BM cells. These data suggest that IRAK1/4 inhibition would be effective at preventing atherosclerosis in individuals with CH carrying ASXL1 mutations. Thus, a tailored treatment targeting specific inflammatory molecules according to driver CH mutations could be a promising strategy to improve the long-term clinical outcomes of CH carriers.

## Acknowledgement

We thank the FACS Core, the Mouse Core and the Pathology Core Laboratories at the Institute of Medical Science, the University of Tokyo. We also thank Daisuke Oikawa, Fuminori Tokunaga, and Mizuki Yamamoto for the plasmids, and Yunong Wang for supporting the analysis of the atherosclerotic lesions. This work was supported by Grant-in-Aid for Scientific Research (A) (No. 20H00537) (to T.K.), Grant-in-Aid for Scientific Research on Innovative Areas (No. 19H04756) (to T.K.), The Tokyo Biochemical Research Foundation (to T.K.), Grant-in-Aid for Scientific Research (B) (No. 19H03685) (to S.G.), The Japan Foundation for Aging and Health (to S.G.), and Koyanagi Foundation (to S.G.).

## Author contributions

N.S. designed and performed most of the experiments, analysed and interpreted the data, and wrote the manuscript. S.G. conceived the project, designed the experiments, interpreted the data, and wrote the manuscript. Y-H.C., T.F., S.A., S.S., X.L., T.Y., R.T., Y.T. and K.Y. assisted with the experiments. T.S. and K.M. provided the BaκB cells and advised on the data interpretation. M.S. and H.S. provided the *Ldlr*^-/-^ mice and advised on the analysis of the atherosclerotic lesions. T.D. and Y.O. assisted in the analysis and staining of the atherosclerotic lesions. S.K. and A.I. performed the RNA-seq experiments. C.L. and E.M. provided the IRAK1/4 inhibitors R191 and R221. J-i.I. advised on the analysis of the TLR-NF-κB pathway. H.H. generated the ASXL1-MT KI mice. T.K. conceived the project, interpreted the data, and wrote the manuscript.

## Conflict-of-interest disclosure

E Masuda and C Lamagna own stock of and are employees of Rigel Pharmaceuticals, Inc. The other authors declare no conflicts of interest.

## Materials and Methods

### Plasmids

To investigate protein-protein interactions in 293T cells, we used 3×FLAG-tagged ASXL1-WT or ASXL1-MT or ASXL1-CT in a pcDNA3.1 vector, and Myc-tagged ASXL1-WT or ASXL1-MT in a pcDNA3.1 vector. We also used FLAG-tagged mTak1 and HA-tagged mTak1 or TAB2 or TAB3 in a pCMV vector, HA-tagged K63Ub and IRAK1 in a pcDNA3.1 vector, FLAG-tagged mTraf6 in a pME18S vector. We amplified the 3×FLAG-tagged C-terminal side of the ASXL1 sequences by PCR and inserted the PCR-amplified sequences into the EcoR1 and Not1 restriction sites of the pMYs-IRES GFP or pMYs-IRES NGFR vector and pcDNA3.1 vector. For NF-κB reporter assay, we used ASXL1-WT, ASXL1-MT and the C-terminal side of ASXL1 in the pMYs-IRES-NGFR vector.

### Mice

C57BL/6J mice were purchased from Japan SLC (Sankyo Labo Service Corporation, Tokyo, Japan). *Ldlr*^-/-^ mice were a gift from the Hitoshi Shimano lab (the University of Tsukuba). *LysM-cre* transgenic mice were a gift from the Takeharu Sakamoto Lab (the University of Tokyo). ASXL1-MT-KI mice were generated in our lab. In brief, a floxed neomycin and stop cassette followed by N-terminal 3xFLAG-tagged mouse ASXL1 mutant (1–1,890) cDNA and internal ribosome entry site (IRES)/enhanced GFP (EGFP) were inserted into the Rosa26 locus by homologous recombination.

Mice carrying the floxed allele (3xFLAG-ASXL1 mutant flox/flox-IRES-EGFP) were crossed with *Vav-cre* transgenic mice or *LysM-cre* transgenic mice. All animal studies were approved by the Animal Care Committee of the Institute of Medical Science, the University of Tokyo (approval number: PA16-54) and conducted in accordance with the Regulation on Animal Experimentation at the University of Tokyo based on International Guiding Principles for Biomedical Research Involving Animals.

### Modeling hyperlipidemia and hypercholesterolemia

To analyze the atherosclerotic lesion in *Ldlr*^-/-^ mice, the mice were started on a high fat, high cholesterol diet (HFHCD) (Research Diets Inc., D12108C, Clinton/Cybulski Diet; 40 kcal%fat, 1.25%cholesterol, no cholic acid) or control diet (Research Diets Inc., 98121701; 10 kcal%fat) after more than four weeks post-transplantation. This hypercholesterolemia-promoting regimen was continued for 12, 14, or 28 weeks, as noted above.

For the analysis of glucose tolerance and the competitive repopulation assay with high-fat diet (HFD), mice were started on a HFD (Research Diets Inc., D12492; 60 kcal%Fat) or control diet (Research Diets Inc., D12450J; 10 kcal%Fat) after four weeks post-transplantation. This hyperlipidemia-promoting regimen was continued for 10 months, as noted above. The IPGTT were performed following 10-week of diet feeding. A bolus of glucose (10 mg/g·body weight) was either injected into the intraperitoneal cavity, and blood was sampled at 0, 15, 30, 60, and 120 min for plasma glucose and insulin analyses after an overnight (16 h) fast.

### IRAK1/4 inhibitor treatment

IRAK1/4 inhibitor (R191 and R221) was kindly provided by Rigel Pharmaceuticals. R221 is a pro-drug of R191 designed for ease of dosing in animal studies. The pro-drug moiety gets cleaved in the lumen of the gastrointestinal tract before R191 is absorbed. Only R191 is found in the systemic circulation. For in vitro treatments, 293T cells, THP-1 cells or RAW264.7 cells were pre-treated with 100 nM R191 1 hour before the lysis in the sample buffer. The *Ldlr*^-/-^ mice transplanted with bone marrow (BM) from ASXL1-MT KI or control mice were fed the HFHCD (Research Diets Inc., D12108C) containing R221 or normal HFHCD after reconstitution for more than 1 month. We performed an analysis of peripheral blood cells 6 weeks after the start of the HFHCD. After 12 weeks of treatment, we analyzed the aortic root using the method as described next.

### Quantitation of atherosclerosis and histological analysis of organs

Serial cryostat sections of the aortic root (6 μm) were cut from OCT-embedded unfixed hearts at the level of the aortic valves and were stored at -80°C until use. Aortic root sections were stained with Oil Red O (ORO) (Muto Pure Chemicals, Cat. No. 4049-2), a lipophilic red dye, to assess plaque accumulation or with Masson’s Trichrome stain (Muto Pure Chemicals, Cat. No. 4020-1, 4025-1) to evaluate sclerosis and fibrosis. Images of the roots were acquired using a NanoZoomer SQ Digital slide scanner (https://www.hamamatsu.com). Quantification of the aortic root lesions was performed using DNP view2 Viewing software (https://www.hamamatsu.com) on 2 or 3 adjacent, ORO-stained cryostat sections. The fibrous cap was defined as an aniline blue-positive structure overlaying the plaque core, with no more than one macrophage foam cell overlying or interpenetrating the cap. Fifteen equally dispersed measurements of the cap thickness were taken for each plaque section. Aortas were cut at ∼5 mm proximally from their main branchpoints and then cleaned of visceral fat deposits in petri dishes filled with cold PBS, opened longitudinally, pinned onto silicon rubber sheets, and then fixed in 10% formalin overnight and stained with Sudan IV (Muto Pure Chemicals, Cat. No. 3307-2). A Canon EOS X80 camera was used to take pictures, and the extent of plaques in the root or aorta or sclerosis in the root was quantified using Adobe Photoshop CC 2019. The total SudanIV staining portion was divided by the total area of the pinned descending aorta to obtain the proportion of the aorta involved in the lesion.

### Cell Culture

293T cells (CRL-11268, ATCC, Manassas, VA, USA) and RAW264.7 cells (TIB-71, ATCC) were cultured in DMEM containing 10% fetal bovine serum (FBS). THP-1 cells (TIB-202, ATCC) were cultured in RPMI-1640 medium supplemented with 10% FBS. Ba/F3 cells with NF-κB-GFP (BaκB cells) were a gift from the Kenske Miyake Lab (the University of Tokyo) and were cultured in RPMI-1640 medium supplemented with 10% FBS containing IL-3. All cell lines used in this study were authenticated by short-tandem repeat analyses and tested for mycoplasma contamination in our laboratory.

Bone-marrow-derived macrophage (BMDMs) were prepared ex vivo from mouse BM cells from ASXL1-MT KI mice or control mice. BM cells were cultured in 10 ml complete DMEM with 10% CMG14-12-conditioned medium at 2 × 10^6^ cells per 10 cm for 3 days, after which cells were cultured for 4 days with another 10 ml fresh medium. After 7 days in culture, adherent cells were >95% CD11b^+^ F4/80^+^, as assessed by flow cytometry.

### Coimmunoprecipitation and western blotting

cells were washed with ice-cold PBS and were lysed with cell lysis buffer (50 mM Tris-HCl, 1 mM EDTA, 150 mM NaCl, 1% TritonX-100, 0.1% SDS, 2 mM Na3VO4, 2mM PMSF, 50mM NaF, 10 μM MG132) supplemented with proteinase inhibitor (cOmplete Mini, Roche, 11836153001). Lysates are immunoprecipitated with anti-FLAG (Sigma, F1804, 1:100) or anti-AKT1/2/3 (Santa Cruz Biotechnology, sc-81434, 1:20) antibody using Dynabeads Protein G (Thermo Fisher Scientific, 10004D) for 10 min at room temperature.

293T cells were transiently transfected with DNA plasmids using polyethylenimine (PEI). The cells were cultured for 48 h after transfection and washed with ice-cold PBS and were lysed in total cell lysis buffer [50 mM Tris-HCl pH7.5, 1 mM EDTA, 150 mM NaCl, 1% Triton X-100, 50 μM NaF, 2 mM PMSF, 2 mM Na3VO4, 0.1% SDS, 10 μM MG132] supplemented with proteinase inhibitor (cOmplete Mini, Roche, 11836153001). For the coimmunoprecipitation (co-IP), Dynabeads Protein G (Invitrogen, 10004D) was pretreated for 10 min at room temperature with anti-FLAG (Sigma-Aldrich, catalog F1804, clone M2), anti-Myc (Santa Cruz Biotechnology, sc-40, clone9E10) or anti-HA (Roche, catalog 11867423001, clone 3F10). Then the sample lysates were incubated with pretreated Dynabeads Protein G for 15 min at room temperature. The precipitates were washed three times with cell lysis buffer, subjected to sodium dodecyl sulfate-polyacrylamide gel electrophoresis (SDS-PAGE) and analyzed by western blotting. Western blotting was performed with anti-DYKDDDDK (Wako, catalog 018-22381, clone 1E6), anti-Myc, anti-HA, anti-phosphorylated TAK1 (CST, catalog 4536S, clone), anti-Lamin B1 (Santa Cruz Biotechnology, sc-374015), anti p38 MAPK (CST, #9212), anto-SAPK/JNK (Cell Signaling Technology, #9252), anti-IκBα (CST, #4814), anti-Phospho-p38 MAPK (Thr180/Tyr182) (CST, #9216), anti-Phospho-SAPK/JNK (Thr183/Tyr185) (CST, #9255), anti-Phospho-IκBα (Ser32/36) (CST, #9246) and anti-GAPDH (CST, #5174). The signals were detected with ECL Western Blotting Substrate (Promega) or ECL Prime Western Blotting Detection Reagent (Cytiva), and immunoreactive bands were visualized using an LAS-4000 Luminescent Image Analyzer (FUJIFILM). Band intensities were analyzed with ImageJ 1.52a software (National Institute of Health, USA).

### Subcellular fractionation

Cells were lysed in hypotonic buffer [10 mM HEPES, 1.5 mM MgCl2, 10 mM KCl, 0.5 mM DDT, and 0.1% TritonX-100 with protease inhibitor cocktail (Sigma)] for 20 min at 4 degrees Celsius and centrifuged (8000 rpm, 5 min), and the supernatant was collected as the cytoplasmic fraction. The insoluble pellet was washed with 100 μL ice-cold PBS once and then lysed in lysis buffer [0.5% NP-40 in TBS with protease inhibitor cocktail (Sigma) and Benzonase (Sigma)] for 30 min at 4 degrees Celsius and centrifuged (14000 rpm, 30 min), and the supernatant was collected as the nuclear fraction.

### Quantification of cytokines

The levels of TNF-α, IL-6, and IL-1β in blood serum were measured with Mouse TNF-alpha Quantikine ELISA Kit (R&D Systems, MTA00B), Mouse IL-6 Quantikine ELISA Kit (R&D Systems, M6000B), and Mouse IL-1β/IL-1F2 Quantikine ELISA Kit (R&D Systems, MLB00C), respectively, according to the manufacturer’s protocol.

### Flow cytometry

Peripheral blood was analyzed using CD11b-PE (eBiosciense, 12-0112-85), CD45R/B220-PE/Cy7 (BioLegend, 103224), CD3-APC (100236), and CD45.2-APC/Cy7 (BioLegend, 109824) antibodies. CD11b-PE/Cy7 (BioLegend, 101216), Ly6G-APC (BioLegend, 127613), Ly6C-Pacific Blue (BioLegend, 128014), CD115-PE (BioLegend, 135505), CD115-Biotin (eBioscience, 13115285), Streptavidin-PerCP-Cy5.5 (eBioscience, 45431782), B220-PE (eBioscience, 12045285), CD3-APC (BioLegend, 100236). Propidium iodide (PI) or 4′,6-diamidino-2-phenylindole (DAPI) was used to exclude dead cells. All data were collected using a FACSVerse with BD FACSuite software or FACSAria with FACSDiva software and were analyzed with FlowJo software.

### NF-κB reporter assay

BaκB cells expressed each TLR (TLR4, TLR5, TLR7). ASXL1-WT and ASXL1-MT were cloned into retroviral vector pMYs-IRES NGFR and transduced into BaκB cells using a retrovirus made from Plat-E packaging cells. The transduced cells were selected using the FACS Aria. BaκB cells were seeded in 96-well culture plates at a density of 10,000 cells per well. The cells were stimulated with Lipid A (E. coli, PEPTIDE INSTITUTE, catalog 24005-s) for TLR4, flagellin (FLA-ST, InvivoGen) for TLR5, and R848 (Invivogen, catalog code tlrl-r848) for TLR7. BaκB cells were harvested 18 h after the stimulation and assayed for NF-κB activity by the mean fluorescence intensity (MFI) using the FACS Verse.

### Immunofluorescence

293T cells transfected with FLAG-tagged ASXL1-WT or ASXL1-MT or empty vector, BMDM cells were subjected to immunofluorescence. The cells were fixed with 4% paraformaldehyde, permeabilized with 0.2% Triton X-100 and blocked with 5% goat serum. The cells were then stained with anti-FLAG antibody, followed by AlexaFluor633-mouse antibody (Thermo Fisher, A21052). The nuclei were visualized with DAPI (Bio Legend). Fluorescence images were analyzed on an EVOS imaging system (ThermoFisher). Images were analyzed with ImageJ.

### Quantitative RT-PCR

Lipopolysaccharide (LPS, E. coli O111; B4, catalog L4391) or RNA was extracted using TRIzol and reverse transcribed using the High Capacity cDNA Reverse Transcription Kit (Applied Biosystems, 4368814) according to the manufacturer’s protocol. cDNA was then subjected to quantitative RT-PCR using a SYBR Select Master Mix (Applied Biosystems). The sequences of the primers used for the qRT-PCR in this study are as follows:

Tnf Forward 5’- TCTTCTCATTCCTGCTTGTGG-3’

Tnf Reverse 5’- GGTCTGGGCCATAGAACTGA-3’

Il-6 Forward 5’- CTTCCATCCAGTTGCCTTCTTG-3’

Il-6 Reverse 5’- AATTAAGCCTCCGACTTGTGAAG-3’

Il-1b Forward 5’-CTGGTGTGTGACGTTCCCATTA-3’

Il-1b Reverse 5’- CCGACAGCACGAGGCTTT-3’

Actb Forward 5’- CTCTGGCTCCTAGCACCATGAAGA-3’

Actb Reverse 5’- GTAAAACGCAGCTCAGTAACAGTCCG-3’

### RNA-seq

We extracted total RNA using the RNeasy Micro Kit (QIAGEN) from CD11b+ F4/80+ aortic plaque macrophages derived from littermate *Ldlr*^-/-^ mice that were transplanted with BM or from littermate ASXL1-WT or ASXL1-MT KI mice (N= 3 for each group). We dissociated the aorta using a dissection microscope and micro-dissecting tools and collected the aortic arch and the ascending, descending, thoracic and abdominal aorta. Whole aortas were rapidly minced in 200 μL aorta dissociation buffer, pipetted with 800 μL aorta dissociation buffer (125 U/ml Collagenase Type XI; Sigma, C7667, 60 U/ml Hyaluronidase type I; Sigma, H3506, 60 U/ml DNase I, 450 U/ml Collagenase Type I; Fujifilm, 037-17603, and 20 mM HEPES in 1x Phosphate Buffered Saline (PBS)) for a total volume of 1 ml, and incubated for 1 h at 37 °C while shaking at 220 rpm. The suspensions were passed through a 40-μm cell strainer into a new tube using the rubber piston of a syringe plunger. The cells were collected by centrifugation and suspended in cold FACS Buffer containing anti-mouse CD16/CD32 Fc receptor blocking antibody and then stained with anti-CD11b antibody and anti-F4/80 andibody. CD11b+ F4/80+ aortic plaque macrophages were sorted with the FACSAria. Total RNA was subjected to reverse transcription and amplification with a SMART-Seq HT Kit for Sequencing (Clontech). After sonication with an ultrasonicator (Covaris), the libraries for RNA sequencing were generated from fragmented DNA with eight cycles of amplification using a NEB Next Ultra DNA Library Prep Kit (New England Biolabs). After the libraries were quantified using the Bioanalyzer (Agilent), the samples were subjected to sequencing with an Hiseq2500 (Illumina), and 61 cycles of the sequencing reaction were performed. TopHat2 (version 2.0.13; with default parameters) and Bowtie2 (version 2.1.0) were used for alignment to the reference mouse genome (mm10 from the University of California, Santa Cruz Genome Browser). Normalization, the removal of batch effects of the count value, and the detection of significant expression differences were done using DESeq2 (version 2.2.1; Love et al., 2014), with raw counts generated from StringTie. The normalized count was log2-transformed and Z-score scaled before performing a principal component and heatmap analyses, except for hierarchical clustering. A GSEA was performed across aging-related genes (Wahlestedt et al., 2013).

### Statistics and reproducibility

Data are expressed as mean ± standard deviation (s.d.) or standard error of mean (s.e.m.). P values were calculated using paired two-tailed Student’s t-test, Unpaired two-tailed Student’s t-test with or without Welch’s correction, or one-way ANOVA with Tukey–Kramer’s post-hoc test. P values < 0.05 were considered significant. All calculations were performed by using GraphPad Prism software. Sample size ‘n’ indicates biological replicates. No randomization or blinding was used and no animals were excluded from the analysis.

