## Supplemental Information for "CH-related Mutant ASXL1 Promotes Atherosclerosis in Mice via Dysregulated Innate Immunity"

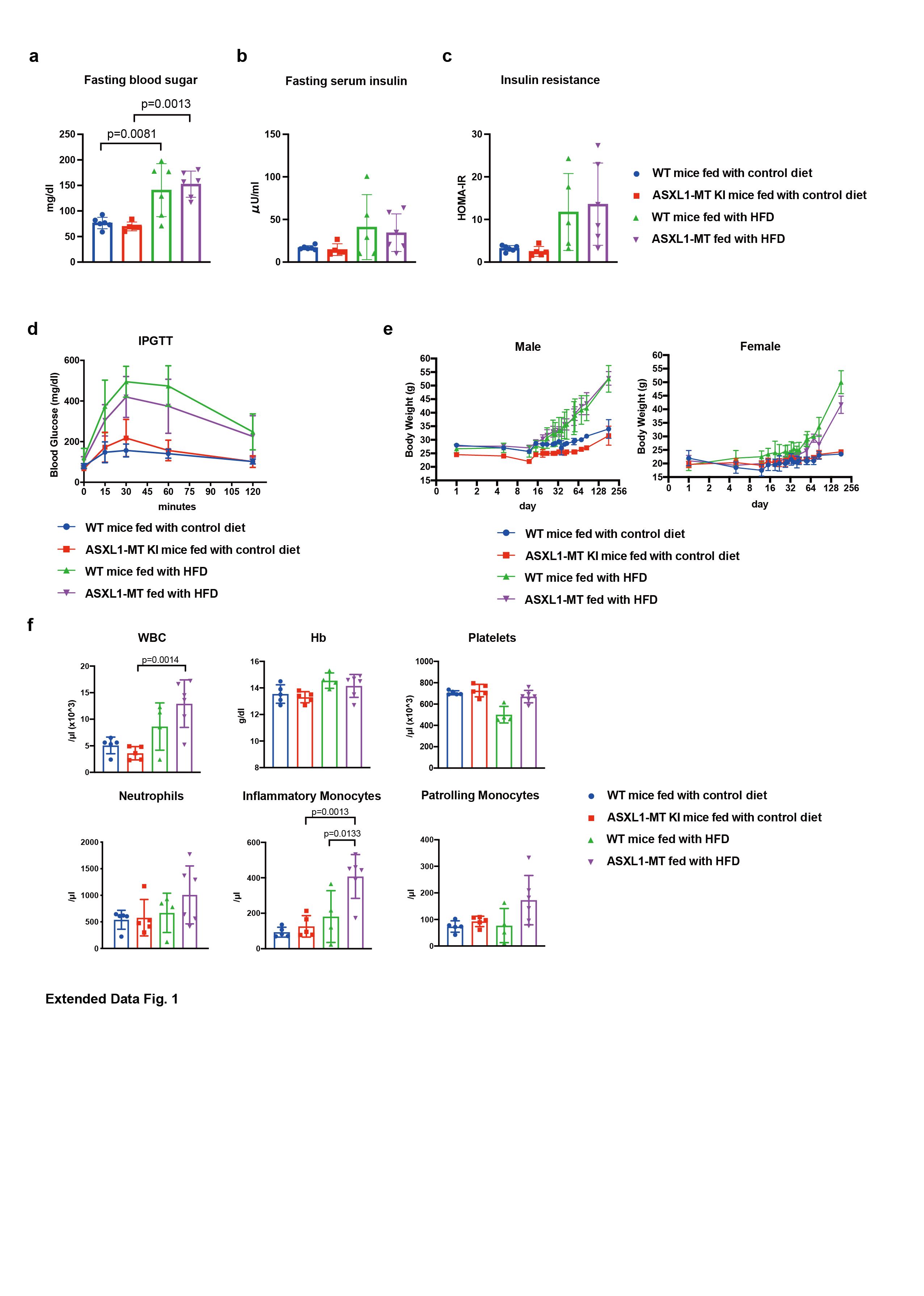

**Extended Data Fig. 1 | Mutant ASXL1 induces expansion of inflammatory monocytes in response to a high-fat diet without any effect on glucose tolerance.**

a-c: The levels of fasting blood sugar (a), fasting serum insulin (b), and homeostasis model assessment as an index of insulin resistance (HOMA-IR) (c) of control or *Vav-cre* ASXL1-MT KI mice fed a control diet or a high-fat diet (HFD) after 10 weeks of feeding.

d: The levels of blood glucose of control or *Vav-cre* ASXL1-MT knock-in (KI) mice fed a control diet or HFD at 10 weeks after feedings in the intraperitoneal glucose tolerance test (IPGTT). The levels of blood glucose were examined at the indicated times after the glucose injection.

e: The body weight of control or *Vav-cre* ASXL1-MT KI mice fed a control diet or HFD at the indicated days after transplantation (left: male, right: female).

f: The absolute number of white blood cells (WBC), platelets, neutrophils, inflammatory monocytes, and patrolling monocytes, and hemoglobin (Hb) in the peripheral blood of control or *Vav-cre* ASXL1-MT KI mice fed a control diet or HFD after 5 months of feeding.

n = 5 (2 female, 3 male), 5 (3 female, 2 male), 4 (2 female, 3 male), 6 (3 female, 3 male) for wild-type (WT) mice fed a control diet, *Vav-cre* ASXL1-MT KI mice fed a control diet, WT mice fed HFD, and *Vav-cre* ASXL1-MT KI mice fed HFD, respectively. Data are the mean ± s.d. Data were assessed by one-way ANOVA with Tukey–Kramer’s post-hoc test.

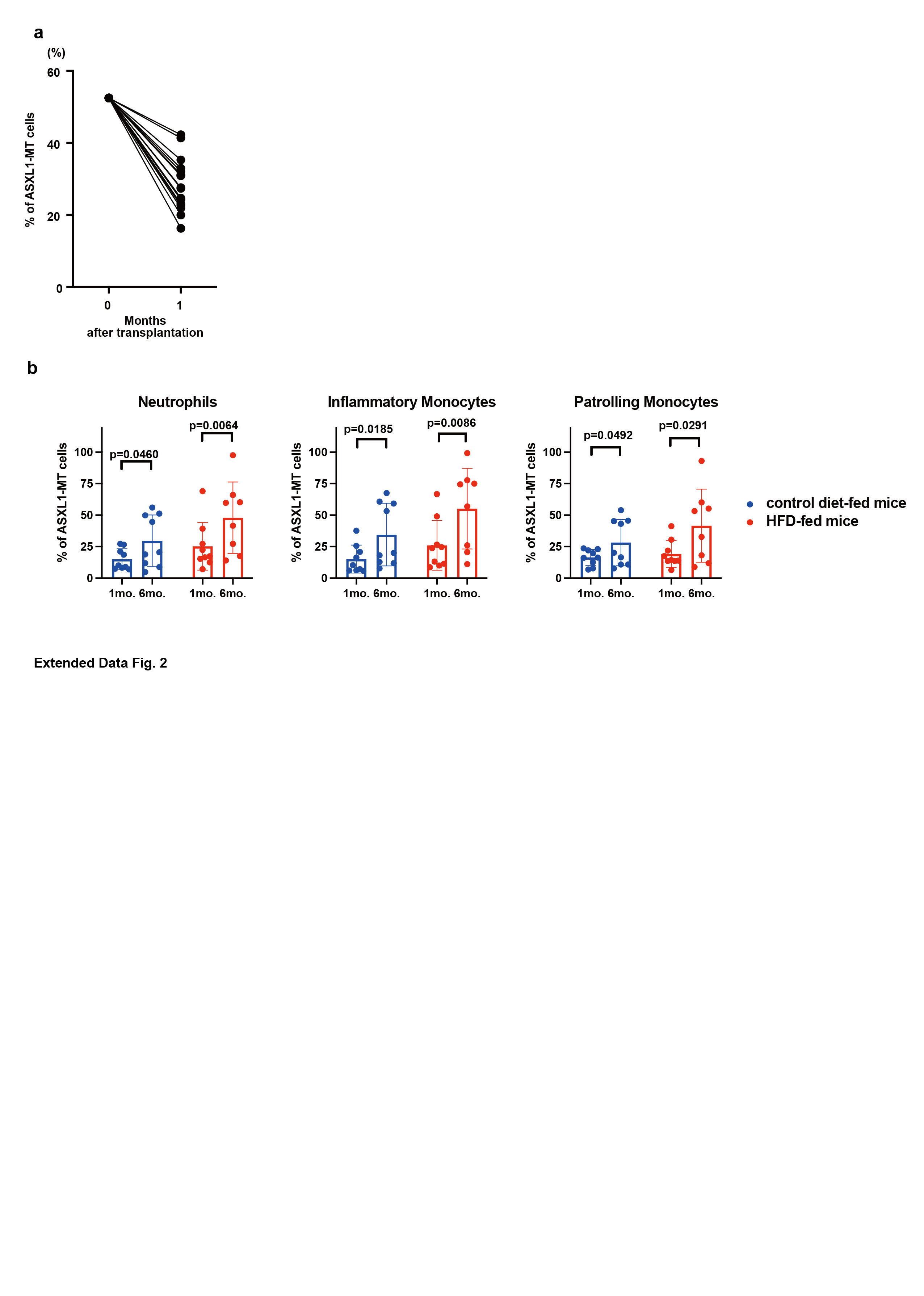

**Extended Data Fig. 2| HFD accelerates the expansion of ASXL1-MT cells in the myeloid population.**

a: The levels of the chimerism of GFP+ ASXL1-MT cells in peripheral blood were analyzed at day 0 and 4 weeks after the transplantation (n = 18).

b: Levels of chimerism of GFP+ ASXL1-MT cells in each myeloid population in PBCs were analyzed at 1 and 6 months after feeding. (n = 9 per group.) Data are presented as the mean ± s.d. Paired t-tests were used for the statistical analysis between each time point (related to Fig. 1b).

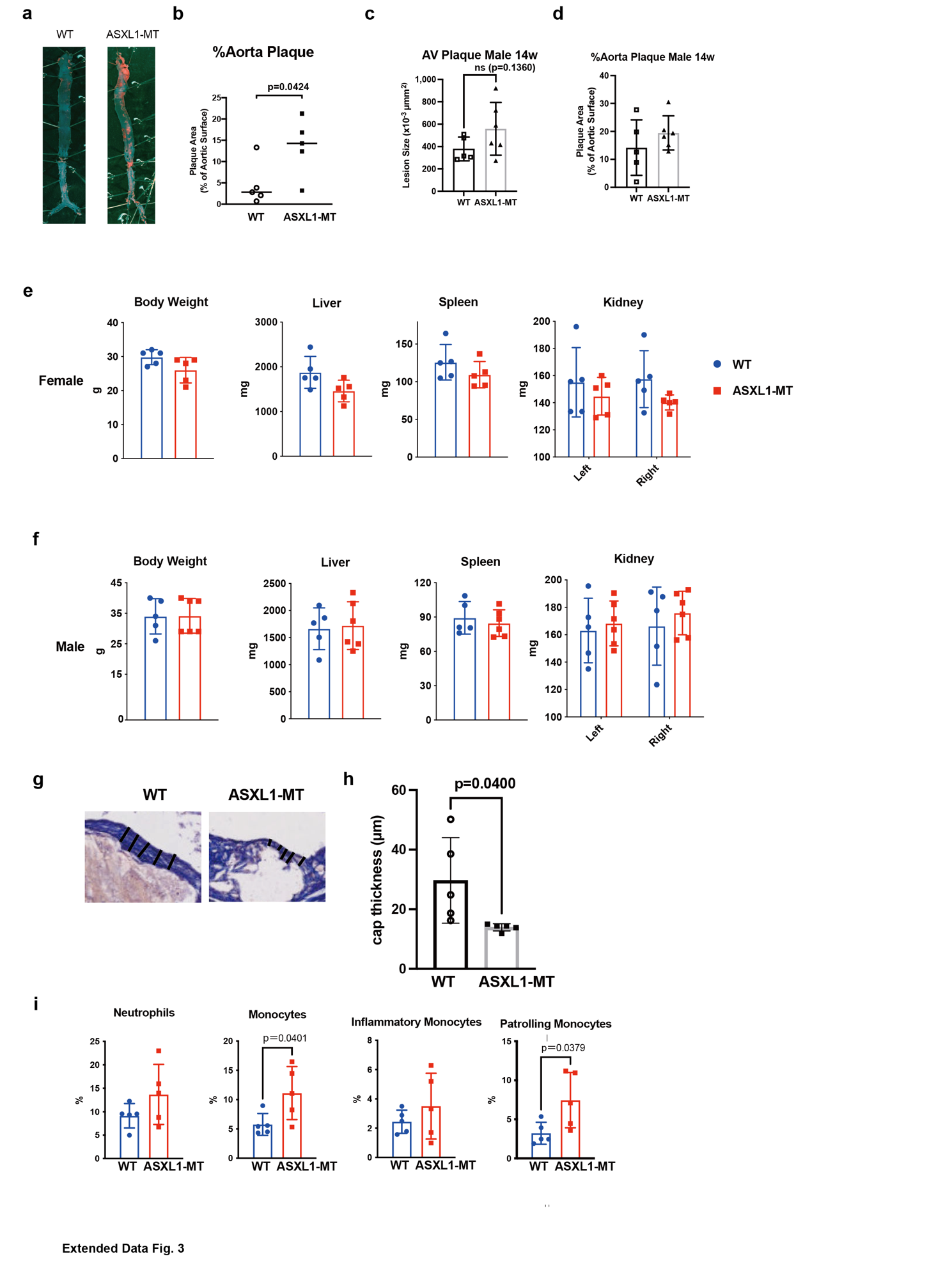

**Extended Data Fig. 3 | Mutant ASXL1-derived hematopoiesis promotes atherosclerosis in *Ldlr*^-/-^ mice.**

a: Representative images of the whole aorta that were stained with Sudan Ⅳ from female
*Ldlr*^-/-^ mice with WT or *Vav-cre* ASXL1-MT KI bone marrow after 12-week HFHCD feeding.

b: Quantification of lesion area in the whole aorta (n = 5 female mice per group).

c, d: Quantification of the lesion area in the aortic root (A) and whole aorta (B) in male *Ldlr*^-/-^ mice with WT or *Vav-cre* ASXL1-MT KI bone marrow after 14 weeks of HFHCD feeding.

(n= 5 WT, 6 *Vav-cre* ASXL1-MT.)

e, f: Bodyweight and major organ weight of female (e) and male (f) *Ldlr*^-/-^ mice transplanted with WT or *Vav-cre* ASXL1-MT KI bone marrow at the analysis of the atherosclerotic lesions. (n =5 WT female, 6 ASXL1-MT female, 5 WT male, 6 ASXL1-MT male.) Data are shown as the mean ± s.d.

g: Representative images of the aortic root lesions that were stained with Masson’s Trichrome stain from female mice *Ldlr*^-/-^ mice with WT or *Vav-cre* ASXL1-MT KI bone marrow after 12-weeks of HFHCD feeding. Black lines indicate cap thickness.

h: Quantification of the cap thickness (n = 5 female mice per group).

i: The frequencies of neutrophils, total monocytes, inflammatory monocytes, and patrolling monocytes in peripheral blood cells in *Ldlr*^-/-^ female mice fed the HFHCD for 12 weeks (n = 9 for each group).

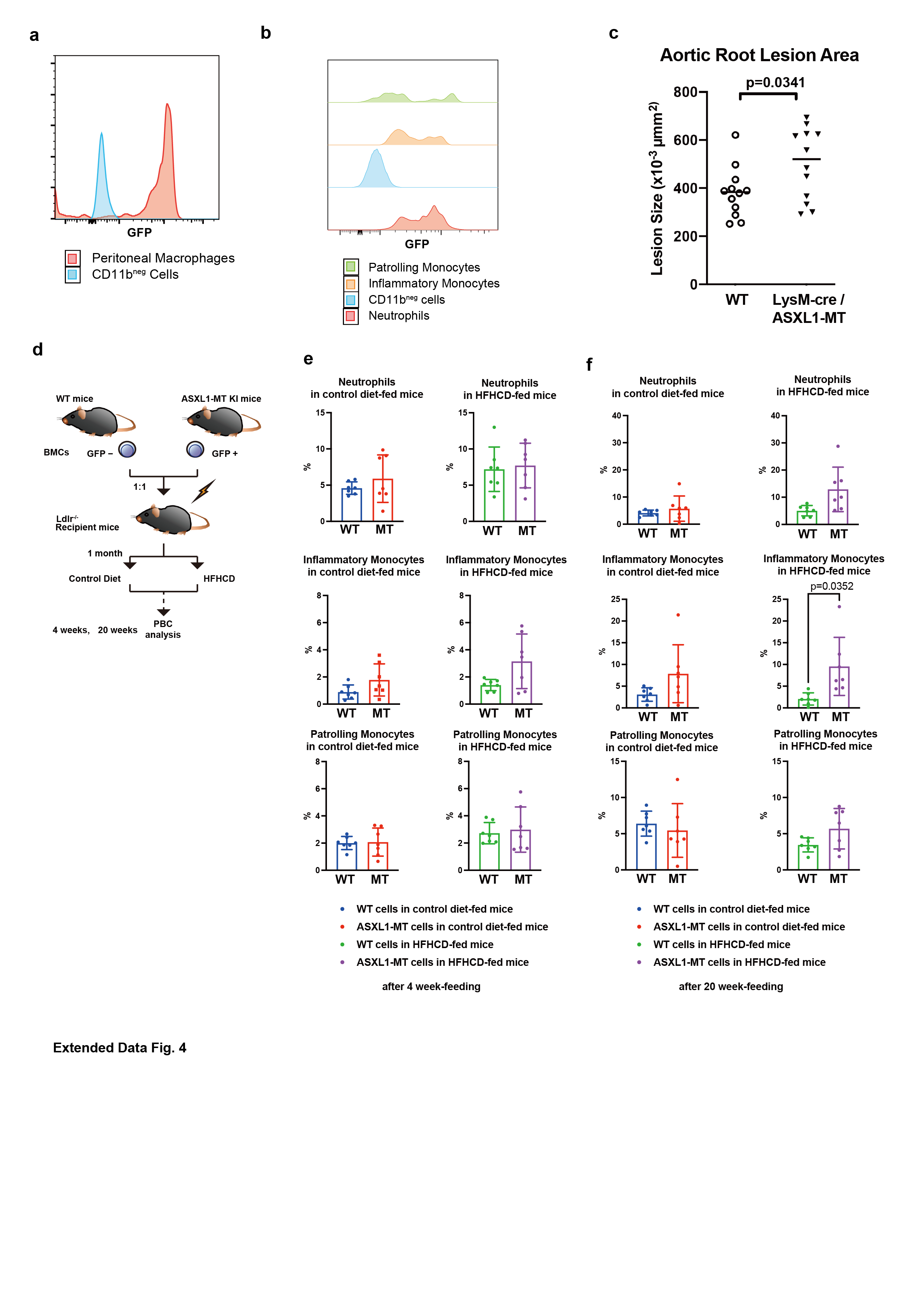

**Extended Data Fig. 4 | Effects of mutant ASXL1-expression on myeloid cells in the atherosclerotic condition.**

a, b: Representative FACS histograms of GFP expression in peritoneal macrophages (a) and peripheral blood myeloid cells (b) from *LysM-cre* ASXL1-MT KI mice. The blue histogram shows the expression of CD11b-negative cells (CD11b^neg^) as a negative control.

c: Quantification of the lesion area in the aortic root from *Ldlr*^-/-^ mice with WT or *LysM-cre*/ ASXL1-MT KI bone marrow after 14-weeks of HFHCD feeding (n = 12 mice per group).
Black horizontal lines represent median values. Unpaired t-tests were used for the statistical analysis.

d: Scheme of the experimental procedures for the competitive transplantation assay in *Ldlr*^-/-^ mice. Briefly, BMCs from *Vav-cre* ASXL1-MT KI mice and littermate control mice were mixed at a 1:1 ratio, followed by transplantation into lethally irradiated *Ldlr*^-/-^ recipient mice. After allowing time for hematopoietic reconstitution, the mice were separated into a HFHCD-fed group and control diet-fed group.

e, f: The frequencies of neutrophils, inflammatory monocytes, and patrolling monocytes in peripheral blood cells derived from GFP^neg^ control bone marrow or GFP^pos^ ASXL1-MT KI bone marrow in *Ldlr*^-/-^ recipient mice that received competitive transplantation and were fed the HFHCD or a control diet for 4 weeks (e) and 20 weeks (f). (n = 7 for each group.) Data were assessed by the paired t-test.

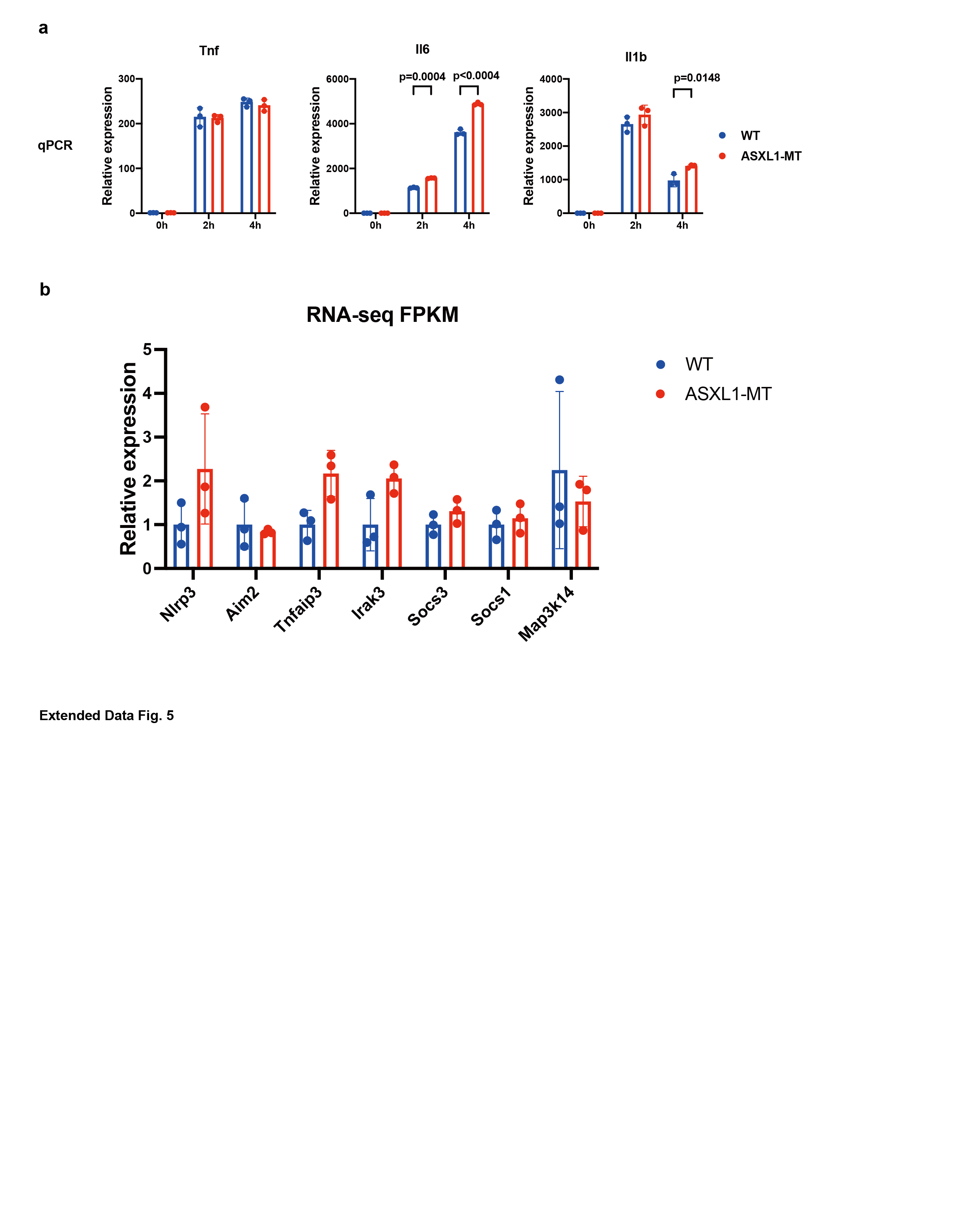

**Extended Data Fig. 5 | Mutant ASXL1-derived macrophages show the inflammatory feature.**

a: Quantitative RT-PCR analysis of the proinflammatory cytokine genes Tnf, Il-6, and Il-1b in wild-type and ASXL1-MT KI BMDMs that treated with lipopolysaccharide (LPS) (1 μg/ml) for the indicated lengths of time. Values and error bars represent the mean ± s.d. of triplicates and are representative of at least two independent experiments.

b: Relative mRNA expression of NF-κB downstream genes in aortic macrophages derived from ASXL1-MT-KI bone marrow or control bone marrow (FPKM).

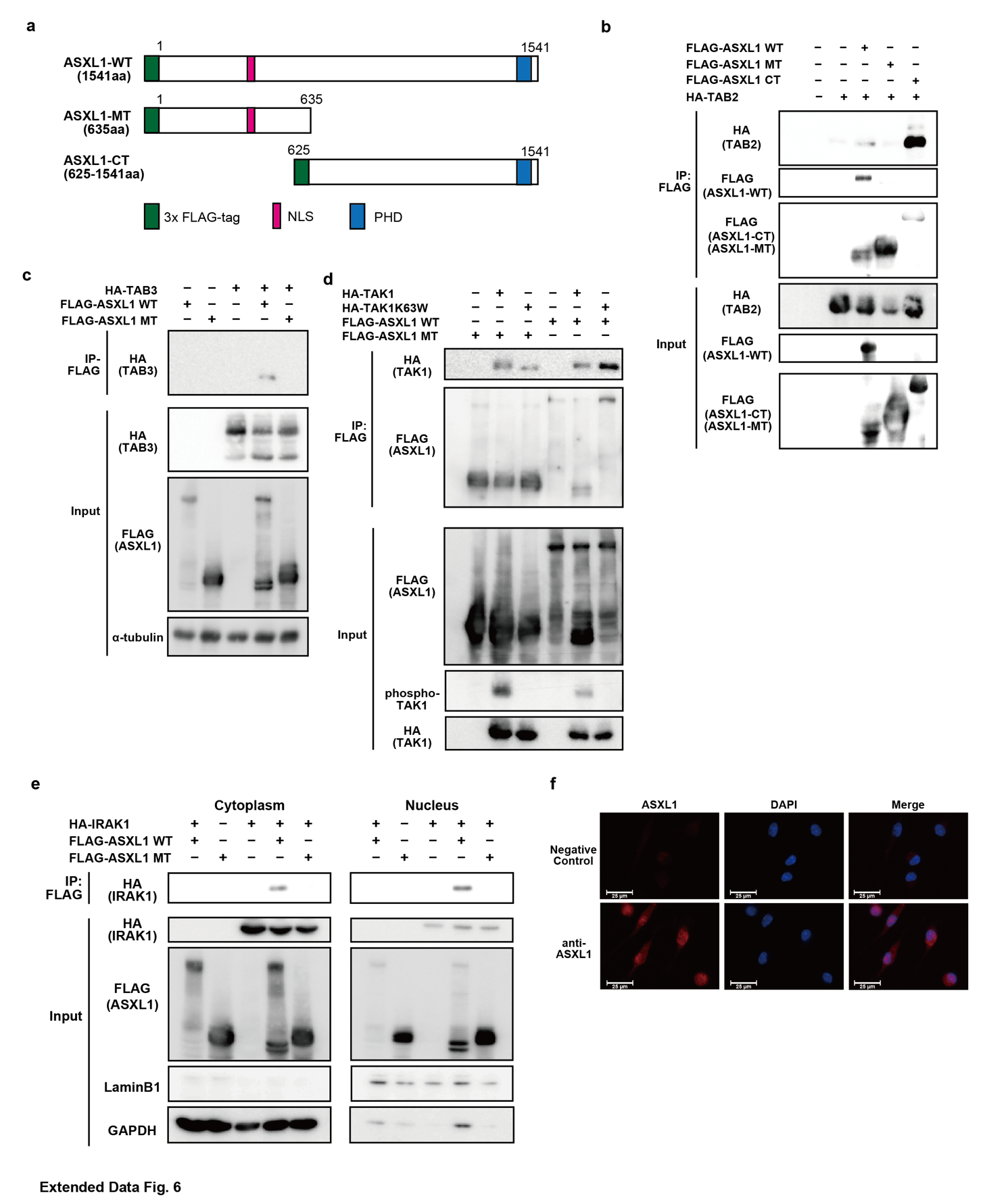

**Extended Data Fig. 6 | ASXL1 interacts with major components of the innate immune pathway in the cytoplasm.**

a: Schematic presentation of the FLAG-tagged ASXL1 constructs: wild-type ASXL1 (ASXL1-WT, 1-1541), C-terminally truncated mutant ASXL1 (ASXL1-MT, 1-645), and C-terminal ASXL1 (ASXL1-CT, 625-1541). PHD: PH domain, NLS: nuclear localization signal.

b, c: 293T cells were transfected with 3xFLAG-ASXL1-WT, 3xFLAG-ASXL1-MT or 3xFLAG-ASXL1-CT expression plasmid together with a plasmid encoding HA-TAB2 (b) or HA-TAB3 (c). Cell lysates were subjected to immunoprecipitation with an anti-FLAG antibody followed by immunoblotting with an anti-HA antibody. Whole-cell lysate (WCL) was prepared and analyzed by immunoblotting with an anti-HA antibody and anti-FLAG antibody.

d: 293T cells transfected with a 3xFLAG-ASXL1-WT or 3xFLAG-ASXL1-MT expression plasmid together with a plasmid encoding HA-tagged TAK1 or HA-tagged catalytically inactive TAK1-K63W were immunoblotted for HA (TAK1 or TAK1-K63W) on immunoprecipitated FLAG-ASXL1. WCL was prepared and analyzed by immunoblotting with anti-HA, anti-FLAG, and anti-phosphorylated TAK1 (phospho-TAK1) antibodies.

e: 293T cells were transfected with a 3xFLAG-ASXL1-WT or 3xFLAG-ASXL1-MT expression plasmid together with a plasmid encoding HA-IRAK1. Nuclear (right) and cytoplasmic (left) protein fractions were separated using hypotonic buffer and subjected to immunoprecipitation with an anti-FLAG antibody followed by immunoblotting with an anti-HA antibody.

f: BMDMs from WT mice were fixed and subjected to immunofluorescence staining. ASXL1 (red) is superimposed over nuclei stained with DAPI (blue). Representative images for negative control (upper) and ASXL1 (lower) are shown. Scale bars: 25 μm. Images were captured by EVOS.

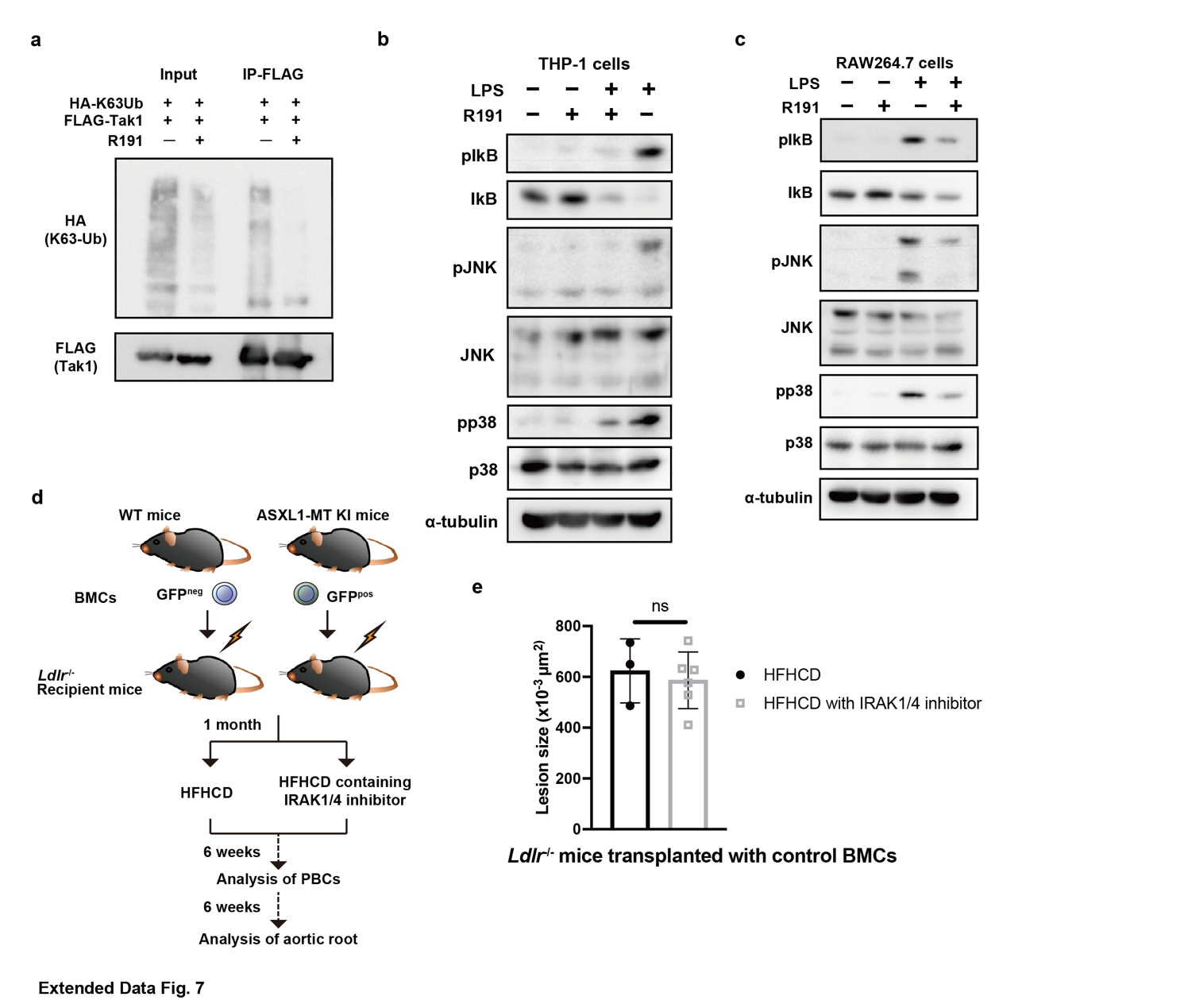

**Extended Data Fig. 7 | Inhibition of IRAK1/4 attenuates activation of downstream TAK1 and ASXL1-MT-driven atherosclerosis.**

a: 293T cells were transfected with FLAG-TAK1 and HA-K63Ub and pretreated with IRAK1/4 inhibitor R191 (100 nM) for 1 h. The K63-linked polyubiquitination of TAK1 was assessed by immunoblotting for HA (K63Ub) on immunoprecipitated FLAG-TAK1.

b, c: THP-1 cells (b) and RAW264.7 cells (c) were pretreated with IRAK1/4 inhibitor R191 (100 nM) for 30 min and stimulated with 1 μg/ml LPS for 30 minutes to evaluate the phosphorylation of IκB, JNK, and p38 MAPK.

d: Outline of the bone marrow transplantations using WT or *Vav-cre* ASXL1-MT KI mice into *Ldlr*^-/-^ recipient mice followed by IRAK1/4 inhibitor (R221) treatment. One-month post-transplantation, the recipient mice were separated into the high fat/high cholesterol diet (HFHCD) group and R221-containing HFHCD group. Peripheral blood cells (PBCs) were analyzed after 6 weeks of feeding. Atherosclerotic lesions were analyzed after 12 weeks of feeding.

e: Quantification of the lesion areas stained with ORO from female *Ldlr*^-/-^ mice that were transplanted with normal murine bone marrow after feeding on a high fat/high cholesterol diet (HFHCD) or IRAK1/4 inhibitor R221-containing HFHCD. (n= 3 mice fed HFHCD, 6 mice fed R221-containing HFHCD). Data are presented as the mean ± s.d. Unpaired t-tests with Welch’s correction were used for the statistical analysis.

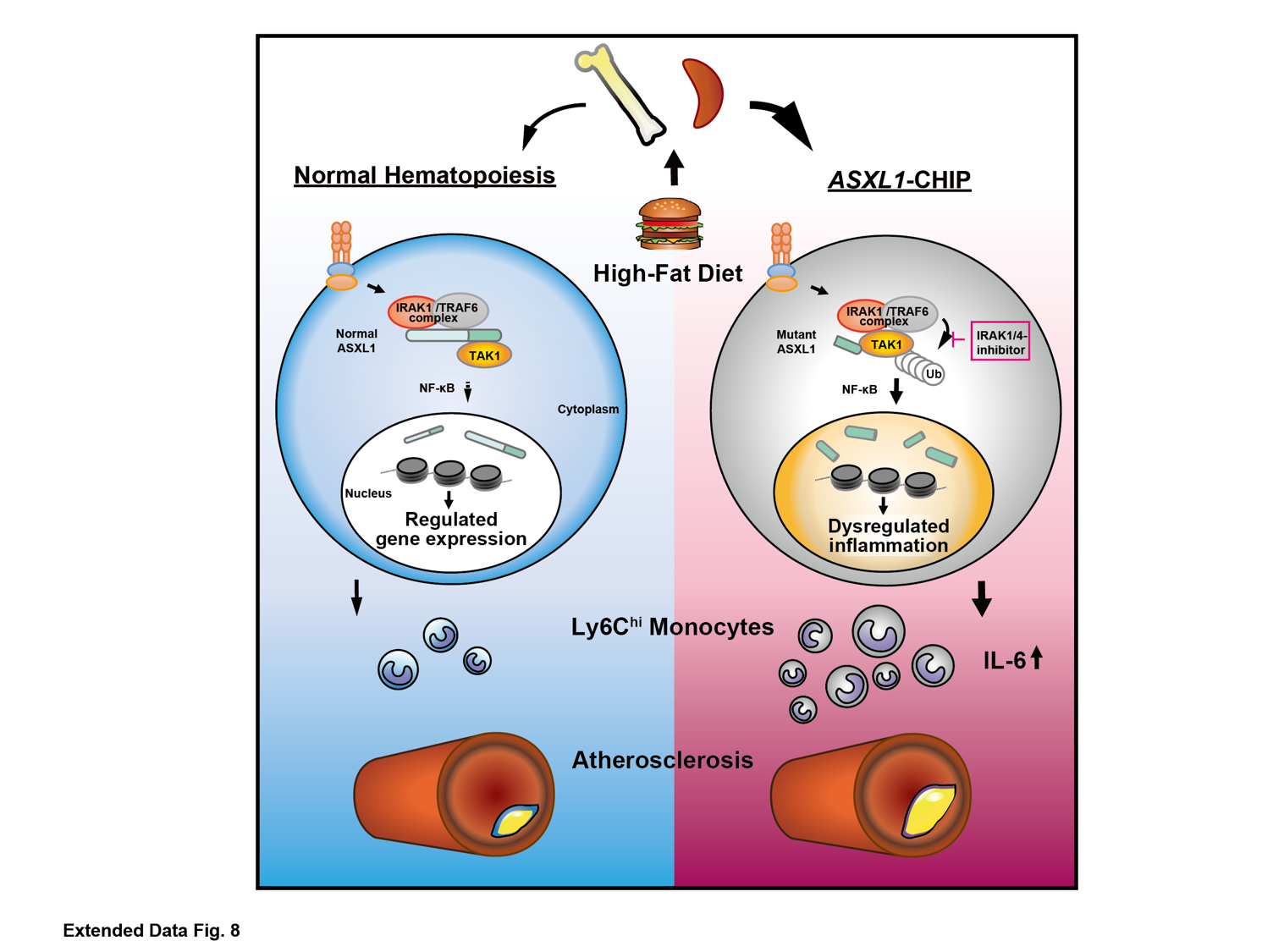

**Extended Data Fig. 8 |** Overall summary scheme.

p=0.0065

p=0.0081

ns

| n= | Diet | Transplanted BMCs | Aortic root  (Mean) | s.d. | 95% CI |
| --- | --- | --- | --- | --- | --- |
| 13 | HFHCD | wild-type | 496.3 | 134.8 | 414.9-577.8 |
| 15 | HFHCD | ASXL1-MT | 639.7 | 121.6 | 572.4-707 |
| 6 | HFHCD containing IRAK1/4 inhibitor | wild-type | 586.7 | 111.7 | 469.5-703.9 |
| 7 | HFHCD containing IRAK1/4 inhibitor | ASXL1-MT | 503.2 | 86.46 | 423.2-583.1 |

**Extended Data Table 1. |**

Total parameters of plaque lesion size in the aortic root of *Ldlr*^-/-^ mice in this study. The number of samples, the kind of diet, the transplanted bone marrow (BM), mean of lesion size in aortic root, standard deviation (s.d.) and 95% confidence interval (CI) are shown. The p values were calculated using unpaired two-tailed Student’s t-test with Welch’s correction.
